# Temporally-segregated dual functions for Gfi1 in the development of retinal direction-selectivity

**DOI:** 10.1101/2025.06.03.657700

**Authors:** Karina Chaudhari, Luis E. Guzman-Clavel, Alex L. Kolodkin

## Abstract

There is great diversity in retinal ganglion cells subtypes in the mouse retina, but little is known about the molecular factors required to generate this subtype diversity. Here, we identify the transcription factor Gfi1 as a conserved driver of differentiation specifically in two downward-tuned direction selective ganglion cells (DSGCs). Further, we describe a post-differentiation role for Gfi1 in regulating dendritic development of Down ON-type DSGCs crucial for their ability to detect downward motion. These results define novel temporally-segregated dual functions for Gfi1 in the development of retinal direction-selective circuits, and they provide a framework for understanding fundamental mechanisms underlying direction-selectivity in the retina.

## Introduction

Direction-selective (DS) neural circuits in the retina are specialized for motion detection in the visual scene. In addition to object tracking, DS circuits also perform the critical function of gaze stabilization in response to global visual motion. These gaze stabilization circuits, termed ON direction-selective circuits, serve as classic models for neural circuit analysis since they control a readily quantifiable reflexive behavior and their isolation from other visual system circuitry renders them easily amenable to manipulation[1, 2]. Within these circuits, the primary detectors of motion are ON DS ganglion cells (oDSGCs) that are tuned to selectively respond to motion in a single direction[3, 4]. oDSGCs are tuned to slow motion and primarily consist of three subtypes responding to up, down or forward motion[4, 5]. Recently, a rare fourth subtype was identified that responds to backward motion[6]. Together with their central midbrain targets in the accessory optic system (AOS), oDSGCs stabilize images on the retina by driving the optokinetic reflex (OKR)[1, 7, 8]. The OKR is an oculomotor behavior that drives compensatory eye tracking movements in response to slow global motion to reduce retinal slip. Despite the importance of retinal direction-selective circuits, the mechanisms underlying the specification of oDSGCs and the molecular factors essential to establish directional tuning remain elusive. Here, we identify a dual role for the transcription factor Growth Factor Independent 1 (Gfi1) in both the specification of Down-oDSGCs (D-oDSGCs) and their responses to downward motion, two essential functions that are segregated across developmental time.

Gfi1 is a zinc-finger transcription factor that was first identified in T cells[9]. Gfi1 has been shown to be important for fate specification in the hematopoietic system[10–14], the inner ear[15], and the intestinal epithelium[16, 17]. In *Drosophila*, the invertebrate Gfi1 homolog senseless is essential for the specification of sensory neurons, including R8 photoreceptors[18, 19]. While canonically known as a transcriptional repressor, Gfi1 along with Atoh1 was also found to function in activation of genes in hair cells [20]. Adding further complexity to roles played by Gfi1 in development, recent work demonstrates that Gfi1 functions to antagonize p53-mediated apoptosis, suggesting that reduced levels of Gfi1 promote worse outcomes in human myeloid leukemia than a complete absence of the protein[21]. Much remains to be understood about the mechanisms of Gfi1 function in different tissues and its critical downstream targets, especially given the clinical importance of Gfi1 in multiple disease states.

Here, we investigate how Gfi1 guides cell-type determination in the mouse retina. We show that Gfi1 is critical for the specification of D-oDSGCs, and identify a novel post-differentiation role for Gfi1 in the development of D-oDSGC responses to downward motion. Further, we find conservation of Gfi1 function in the specification of another distinct DS RGC subtype: the F-mini ONs. Together, these results highlight the multifunctional nature of Gfi1 in the development of DS circuits in the retina, and they reveal a novel role for Gfi1 in the establishment of downward motion encoding.

## Results

### The transcription factor *Gfi1* is selectively expressed in D-oDSGCs within the Accessary Optic System

To identify the mechanisms controlling oDSGC subtype specification and directional tuning, we analyzed the transcriptomic profiles of closely-related D-oDSGCs and Up-oDSGCs (U-oDSGCs) in our previously published dataset of vertically tuned oDSGCs[22]. We noted that the transcription factor Gfi1 shows select expression in D-oDSGCs, with higher expression levels during the early stages of D-oDSGC development (Figures S1A and S1B). The expression levels of *Gfi1* in D-oDSGCs are comparable to those of the transcription factor *Tbx5* in U-oDSGCs, a factor that we have previously found to be essential for U-oDSGC formation[22] (Figures S1A–C). Given the function of Gfi1 in the fate specification of other tissue systems[10–17], we asked whether Gfi1 is essential for D-oDSGC development.

First, to confirm *Gfi1* selective expression in D-oDSGCs within the AOS, we obtained a previously-generated *Gfi1^Cre^* mouse line[23] and crossed it to the *Ai2* and *Ai14* Cre-dependent reporter lines. Wholemount retinas collected from *Gfi1^Cre/+^; Ai2* mice at postnatal day 6 (P6) showed sparse Gfi1 expression. All Gfi1^+^ cells co-label with Rbpms, a pan-RGC marker, indicating that *Gfi1* is expressed exclusively in RGCs at this developmental stage at a density of ∼500 cells/mm^2^ (Figures 1B–1D). Gfi1^+^ RGCs do not exhibit mosaic spacing (Figure 1B), suggesting that Gfi1 is expressed in more than one subset of RGCs[24]. To determine if *Gfi1* is expressed in D-oDSGCs, we performed *in situ hybridization* on P4 retina cryosections. We chose to conduct expression analysis at P4 since *Gfi1* expression starts to decline by the end of the first postnatal week (Figure S1B). *Fibcd1* is expressed exclusively in D-oDSGCs and in subsets of amacrine cells[22], and we detected *Gfi1* expression in *Fibcd1^+^* D-oDSGCs (Figure 1E). Since at P4 *Gfi1* is expressed exclusively in RGCs (Figure S1D), any cell co-expressing *Fibcd1* and *Gfi1* must be a D-oDSGC. We found that ∼15% of *Gfi1*-expressing RGCs co-express *Fibcd1*, consistent with oDSGCs being a relatively rare population in the retina[5, 22, 25, 26] (Figure 1F). We observed no Gfi1 expression in *Tbx5^+^* U-oDSGCs (Figure S1D), nor do P10 *Gfi1^Cre/+^, Ai14, Spig1^GFP^* retinas show any Gfi1^+^ RGCs expressing *Spig1^GFP^*, an U-oDSGC marker[25, 26] (Figure S1E), confirming the absence of *Gfi1* expression by U-oDSGCs. Since oDSGC subyptes innervate distinct AOS nuclei in the midbrain, we examined the innervation patterns of *Gfi1*-expressing RGC subtypes. We analyzed the brains of *Gfi1^Cre/+^, Ai2* and *Gfi1^Cre/+^, Ai14* mice injected intraocularly with the fluorescent dye Cholera toxin subunit B-Alexa 647 (CTB-647), which is transported anterogradely in RGC axons and labels all retinorecipient regions in the brain. Coronal brain sections show strong Gfi1^+^ axonal innervation of the ventral medial terminal nucleus (MTN), the target of D-oDSGCs, but not the dorsal MTN nor the NOT and DTN, targets of U-oDSGCs and Forward-oDSGCs (F-oDSGCs), respectively[5, 25, 26] (Figures 1G, 1H, 1K, 1L and 1N). Taken together, these results show that within the AOS, *Gfi1* is exclusively expressed in D-oDSGCs.

**Figure 1.**
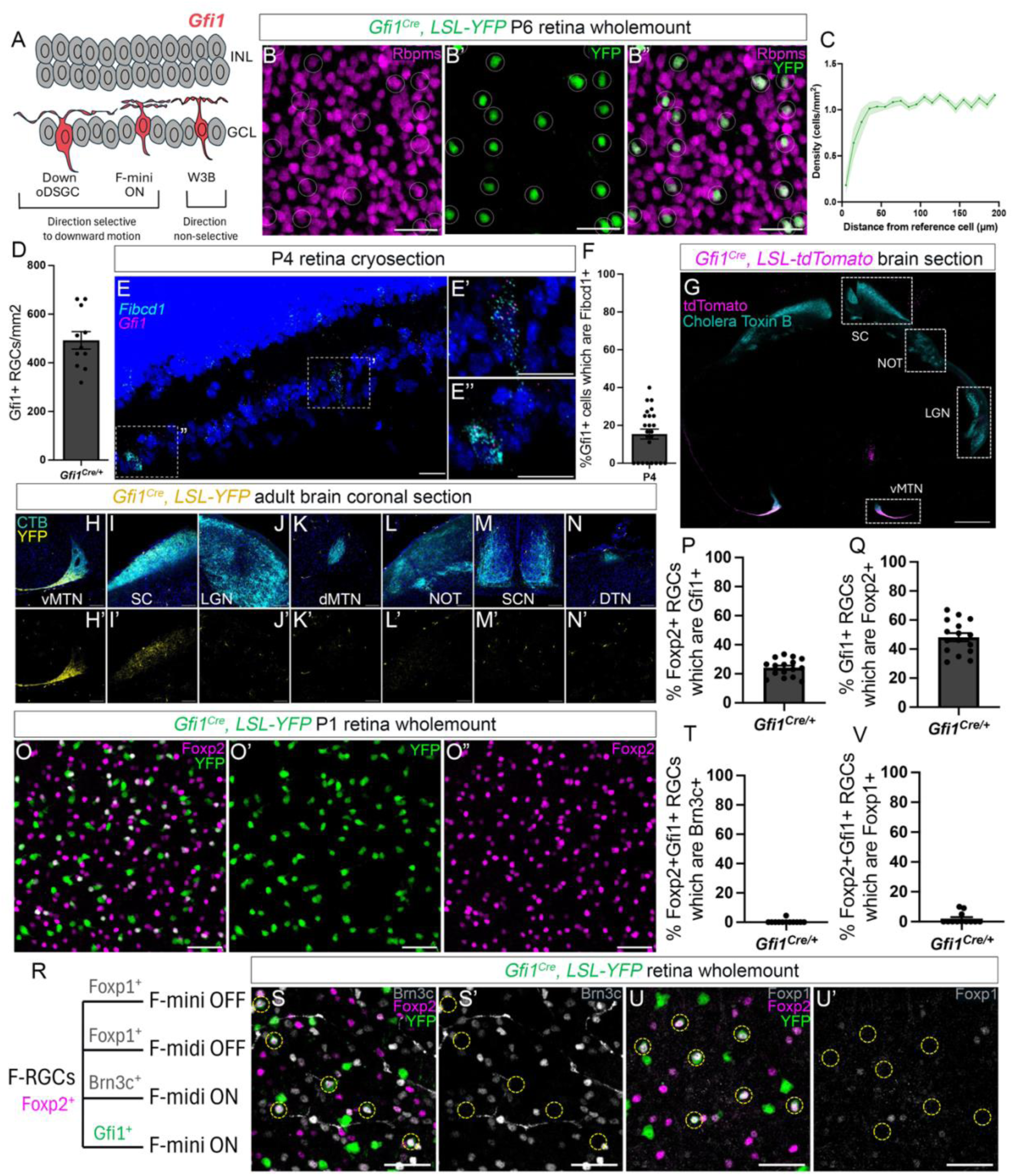
***Gfi1* is expressed in direction-selective D-oDSGC and F-mini ON RGC subsets** (**A**) Schematic depicting *Gfi1* expression in the retina restricted to D-oDSGCs, F-mini-ONs and W3Bs. D-oDSGCs and F-mini ONs are direction-selective subsets tuned primarily to downward motion, while W3Bs are non-direction-selective. (**B**) Wholemount P6 *Gfi1^Cre^, LSL-YFP* retina shows *YFP* expression restricted to Rbpms^+^ RGCs. (**C**) Density recovery profile (DRP) of YFP^+^ RGCs suggests an absence of mosaicism confirming Gfi1 expression in more than one RGC subtype. (**D**) Density of Gfi1-expressing RGCs in the retina. (**E**) *In situ* hybridization in P4 retina cryosection shows *Gfi1* expression in *Fibcd1*^+^ D-oDSGCs. (**E’, E”**) Detailed views of the nuclei indicated in D. (**F**) Quantification of the percentage of *Gfi1^+^* RGCs co-expressing *Fibcd1*. (**G**) Coronal brain section from *Gfi1^Cre^, LSL-tdTomato* adult mouse shows strong innervation of the ventral medial terminal nucleus (MTN) by tdTomato^+^ RGCs. (**H-N**) Detailed views of all retinorecipient nuclei labeled with cholera toxin B (CTB) from *Gfi1^Cre^, LSL-YFP* adult mice confirms YFP^+^ RGC innervation exclusively in the ventral MTN **(H, H’**) and the SC (**I, I’**), and not in the other retinorecipient nuclei (**J-N**). (**O**) Wholemount P1 *Gfi1^Cre^, LSL-YFP* retina shows *YFP* expression in a subset of Foxp2^+^ RGCs. (**P, Q**) These YFP^+^ Foxp2^+^ RGCs account for ∼25% of all Foxp2^+^ RGCs and ∼50% of all Gfi1^+^ RGCs. (**R**) Combinatorial labeling can differentiate between F-RGC subsets. (**S-U**) Wholemount *Gfi1^Cre^, LSL-YFP* retina shows that YFP^+^ Foxp2^+^ RGCs do not express the F-RGC OFF marker Foxp1 or the F-midi ON marker Brn3c, confirming their F-mini ON identity. Data are presented as mean ± SE.

### Gfi1 is also expressed in vertically-preferring direction-selective F-mini ON retinal ganglion cells

In addition to the ventral MTN, we also found Gfi1^+^ axonal innervation in the superior colliculus (SC), reflecting *Gfi1* expression in other non-AOS RGC subsets (Figure 1G, 1I). Querying *Gfi1* expression in existing RGC scRNA-seq databases at P5[27] and P56[28] reveals highly selective expression of *Gfi1* in clusters that include D-oDSGC and additionally, clusters containing W3B and F-mini ON RGCs. While W3Bs are object motion-sensitive but not direction-selective[29], F-mini ONs are one of the four subsets of Foxp2^+^ RGCs (F-RGCs) that are both orientation- and direction-selective, with preferred speeds faster than oDSGCs (∼0.58mm/s vs ∼0.25mm/s)[30]. Analysis of available scRNA-seq databases suggests that Gfi1 is not expressed in the remaining three subsets of F-RGCs: F-midi ONs, F-midi OFFs and F-mini OFFs [27, 28]. Though F-midi ON and OFF subsets are not DS, and the F-mini OFF subtype is directionally tuned to upward motion, the majority of F-mini ONs respond primarily to downward motion[30]. Therefore, F-mini ONs are an interesting RGC subtype for investigating conserved Gfi1-driven mechanisms that establish downward directional tuning. Wholemount retinas collected from *Gfi1^Cre/+^; Ai2* mice at P1 show *Gfi1* expression in approximately 25% of Foxp2^+^ RGCs (Figures 1O and 1P). These Gfi1^+^, Foxp2^+^ RGCs account for roughly half of the RGCs that express *Gfi1* (Figure 1Q).

To classify the *Gfi1*-expressing RGC subtypes, we conducted a morphological assessment by injecting intraocularly a Cre-dependent *AAV9-Brainbow* (*AAV-BbChT*) virus into *Gfi1^Cre^* mice at P0 and visualizing the dendrites of sparsely-labeled RGCs at P15. We found two populations of *Gfi1*-expressing RGCs that do not express Foxp2. One subset had long dendrites stratifying primarily in S4 but with some of these RGCs extending a small fraction of their dendritic arbor into S2, consistent with D-oDSGC morphology[5] (Figure S2A); and another subset had very short dendrites stratifying in S3, a dendritic pattern previously described for W3Bs[29] (Figure S2C). *Gfi1*-expressing RGCs which express Foxp2 showed dendrites within S3 that are partially bistratified (Figure S2B), and, unlike most D-oDSGCs, they almost always showed strongly asymmetric dendritic arbors (Figures S2D and S2E). Their stratification in the region of S3 suggests that these *Gfi1*-expressing F-RGCs are ON and not OFF subtypes since OFFs stratify in S1[30]. We confirmed this is indeed the case by the absence of Foxp1 expression (an F-RGC OFF marker[30], Figure 1R) in Gfi1^+^ Foxp2^+^ RGCs (Figures 2S and 2T). We also excluded *Gfi1* expression in F-midi ONs using Brn3c immunostaining since Brn3c is expressed in F-midi ON RGCs[30] (Figures 2R, 2U and 2V). These data demonstrate conclusively that in addition to D-oDSGCs, *Gfi1* is expressed in W3Bs, which are non-DS, and also in F-mini ONs, another class of DS RGCs that is tuned primarily to downward motion (Figure 1A). Using the *Gfi1^Cre^* line, we observed *Gfi1*-expression in microglia, as has been previously documented with this line[31], and between P6 and P10 we also found a few cells in the inner nuclear layer (INL) that express *Gfi1* (Figures S2F and S2G). Gfi1 can repress its own locus through an autoregulatory feedback loop[11]. Therefore, INL expression of *Gfi1* is likely ectopic expression due to a haploinsufficiency resulting from the Cre knock-in at the *Gfi1* locus[31], since no *Gfi1* expression is noted in any amacrine or bipolar cells in published scRNA-seq databases[32, 33]. We confirmed that *Gfi1* expression is indeed restricted to RGCs using a tamoxifen-inducible *Gfi1^CreERT2^* line[34] crossed to *Ai2*, in which daily tamoxifen injections from P5 to P13 revealed *Gfi1* expression only in some RGCs that are still actively expressing *Gfi1*, and we confirmed that there is no *Gfi1* expression in the INL (Figures S2H and S2I). Taken together, these results show that *Gfi1* expression in the retina is restricted to three RGC subtypes, namely direction-selective D-oDSGCs and F-mini ONs, and W3Bs, which are not DS.

**Figure 2.**
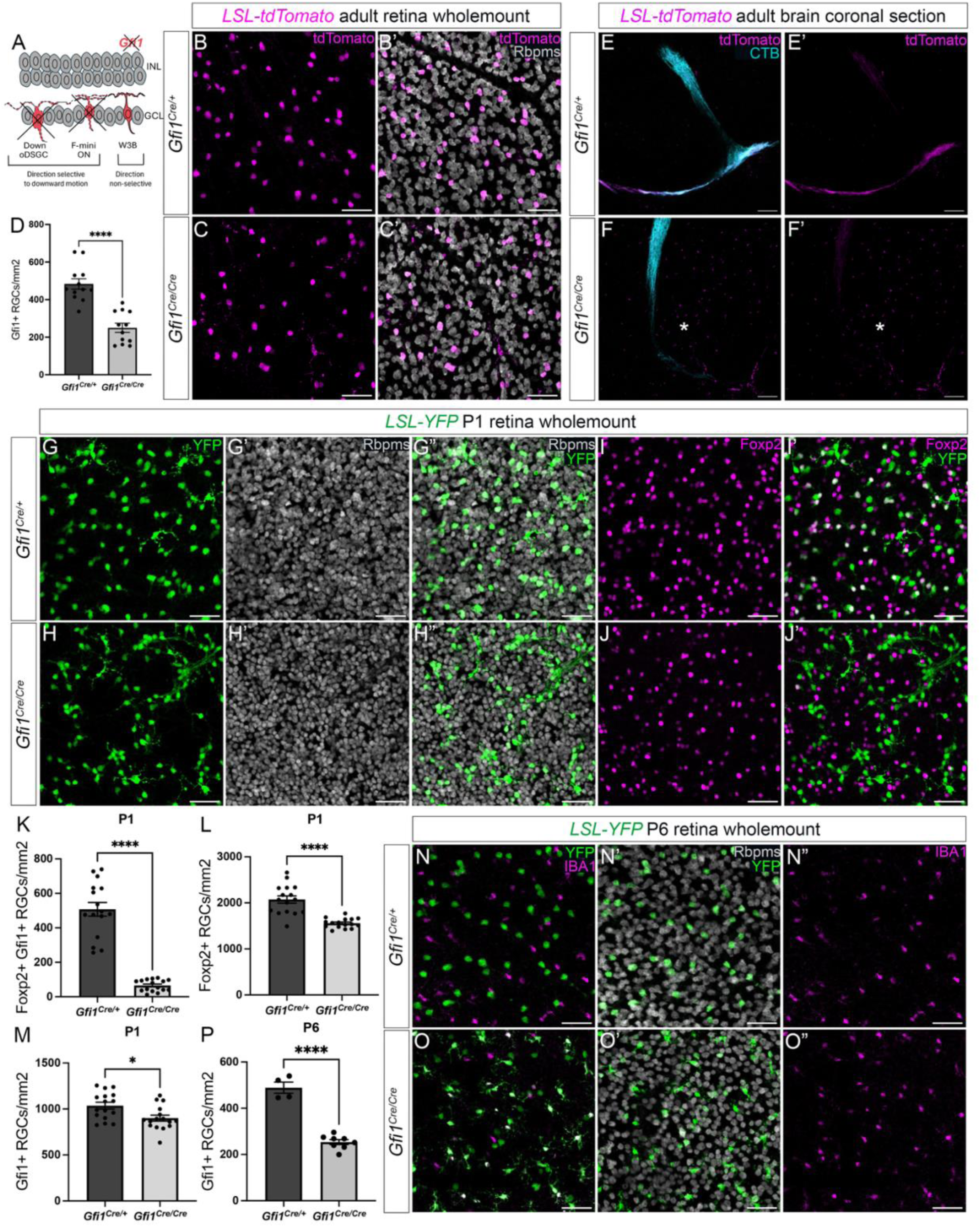
***Gfi1* is essential for the specification of D-oDSGCs and F-mini ONs** (**A**) Schematic depicting a selective requirement for Gfi1 in the specification of the direction-selective D-oDSGC and F-mini-ON subsets. (**B-D**) Adult *Gfi1^Cre/Cre^, LSL-tdTomato* retinas show a loss of ∼50% of tdTomato^+^ RGCs. (**E-F**) *Gfi1^Cre/Cre^, LSL-tdTomato* mice reveal a loss of D-oDSGCs, confirmed by the lack of ventral MTN innervation (indicated by asterisks). (**G-J**) P1 retinas from *Gfi1^Cre/Cre^, LSL-YFP* mice show a ∼25% reduction in Foxp2^+^ RGCs (**L**), reflecting the loss of YFP^+^ Foxp2^+^ F-mini ONs (**I-K**), but only a small change in YFP^+^ RGCs (**M**), suggesting the presence of improperly specified RGCs. Note the excessive clumping of YFP^+^ RGCs (**H**). (**N-P**) By P6, the ∼50% reduction in YFP^+^ RGCs is apparent, suggesting a loss of these improperly specified RGCs between P1 and P6. Data are presented as mean ± SE. *p < 0.05, ****p < 0.0001.

### Gfi1 is essential for the specification of D-oDSGCs and F-mini ONs

The *Gfi1^Cre^* mouse line is a knock-in line in which *Cre* is inserted into the *Gfi1* locus and disrupts *Gfi1* function[23], allowing us to use homozygous *Gfi1^Cre/Cre^* mice to simultaneously remove *Gfi1* and track *Gfi1*-expressing RGCs at different developmental timepoints. Homozygous *Gfi1^Cre/Cre^; Ai14* adult retinas show a loss of approximately half the population of *Gfi1*-expressing RGCs (Figures 2B–2D). We also observed loss of Gfi1^+^ RGC axon innervation of the ventral MTN in *Gfi1^Cre/Cre^; Ai14* adult brains, suggesting that D-oDSGCs are lost upon Gfi1 removal (Figures 2E and 2F). To ensure that loss of ventral MTN innervation is a result of a loss of D-oDSGCs and not simply a loss of *Gfi1* expression by D-oDSGCs, we injected CTB-647 intraocularly into these mice and found that the entire ventral MTN tract was indeed absent, confirming loss of D-oDSGCs (Figures 2E and 2F). We observed no difference in RGC projections to the dorsal MTN, the NOT or DTN (Figures S3A–S3F), or the number of *Spig1^GFP^* U-oDSGCs (Figures S3G–S3I), indicating that this deficit is specific to the D-oDSGCs within the AOS.

To determine whether F-mini ONs are also lost upon Gfi1 removal, we immunostained *Gfi1^Cre/Cre^; Ai12* retinas for Foxp2 at P1. We found a substantial loss of Gfi1^+^ Foxp2^+^ RGCs, reflecting loss of Gfi1^+^ F-mini ONs (Figures 2I’, 2J’ and 2K). To confirm that the absence of Gfi1^+^ Foxp2^+^ cells is due to F-mini ONs no longer being specified, and not merely because F-mini ONs no longer express Gfi1, we assessed total F-RGC numbers. We found that the F-RGC population was reduced by ∼25%, confirming that Gfi1^+^ F-mini ONs are indeed no longer being specified (Figures 2I, 2J and 2L). Despite the lack of identity-specific markers for F-mini ONs at P1 and a 50% loss of Gfi1^+^ RGCs in adults, we did not see the total *Gfi1*-expressing RGC population reduced by half at P1 (Figure 2G, 2H and 2M); we found only a small reduction in the number of Gfi1^+^ RGCs at this very early postnatal time point, suggesting the presence of improperly specified Gfi1^+^ RGCs that are formed initially but are then lost over time. Many of these Gfi1^+^ RGCs are present in aggregates at P1 (Figure 2H) and are lost by P6 (Figures 2N, 2O and 2P). We cannot conclusively show that the Gf1^+^ RGCs that persist are W3Bs due to the lack of specific markers for this RGC subtype; however, our results suggest that Gfi1 removal selectively affects the generation of DS D-oDSGCs and F-mini ON subtypes, while the non-DS W3B subtype is likely unaffected (Figure 2A). Collectively, our results identify a vital role for Gfi1 in the generation of D-oDSGCs and F-mini ONs. Removal of *Gfi1* results in improper specification of these subtypes confirmed by the lack of identity-specific markers, followed by their loss soon after birth.

### Gfi1 is necessary for the downward optokinetic reflex

Previously, we found that a selective loss of U-oDSGCs upon *Tbx5* removal results in a loss of both the upward and downward OKR, despite the preservation of D-oDSGCs[22]. Therefore, we investigated whether the selective loss of D-oDSGCs also affects both the upward and downward OKR. To assess the behavioral effects of Gfi1 removal on the OKR, we turned to *Gfi1* conditional knockout mice[35] since *Gfi1* global mutants are deaf and show a complete loss of inner ear hair cells in both the vestibule and the cochlea[15]; these resultant deficits in their vestibulo-ocular response will confound any OKR measurements. We used *Chx10^Cre^*, a well-characterized line in which *Cre* is expressed in retinal progenitor cells starting at ∼E10.5[36], to achieve a retina-specific knockout of *Gfi1*. We then measured the OKR of these mice, which is a direct behavioral readout of oDSGC and AOS function[1, 5]. In response to continuously-rotating stimuli, *Chx10^Cre^; Gfi1^F/F^* mice showed a selective deficit in the downward OKR, while the upward and horizontal OKRs remained unchanged (Figures 3A–3F). The deficits were specific to OKR since voluntary vertical saccadic movements, including downward displacement of the eye, were normal (Figure 3G). Additionally, in response to oscillating stimuli, these mice showed an increased horizontal gain in response to vertically-moving stimuli (Figures 3H and 3I) – a phenomenon termed “cross-coupling”, which suggests failure of the ventral MTN to suppress the NOT, which is critical for preventing horizontal OKR responses during vertical OKR assessment[22, 37–39]. Altogether, these results demonstrate that Gfi1 functions autonomously in the retina to specifically regulate the downward OKR.

**Figure 3.**
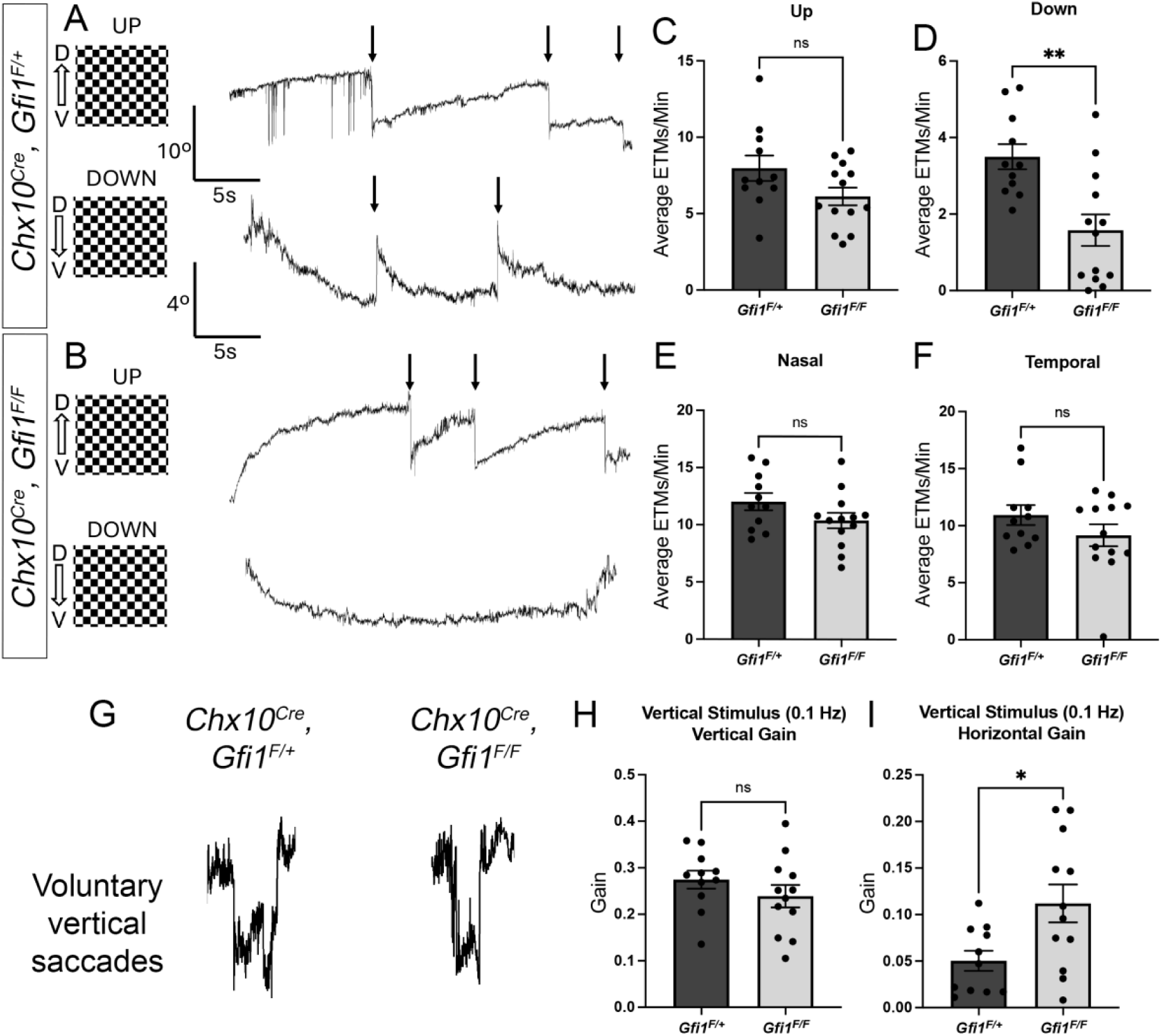
***Gfi1* conditional mutants show selective deficits in downward motion detection** (**A-F**) Optokinetic reflex (OKR) results in adult *Chx10^Cre^, Gfi1^F/+^* and *Chx10^Cre^, Gfi1^F/F^* mice in response to upward (**C**), downward (**D**), forward (**E**), and backward (**F**) continuous motion, quantified as eye-tracking movements (ETMs) per minute (arrows in **A** and **B**). *Chx10^Cre^, Gfi1^F/F^* mice show a selective deficit in downward OKR (**B, D**). (**G**) *Chx10^Cre^, Gfi1^F/+^* and *Chx10^Cre^, Gfi1^F/F^* mice both performed voluntary vertical saccades. (**H, I**) *Chx10^Cre^, Gfi1^F/F^* mice show no change in vertical gain (**H**) but an enhanced horizontal gain (**I**) in response to vertical sinusoidal stimuli. Data are presented as mean ± SE. *p < 0.05, **p < 0.01.

### A temporally-segregated dual-function of Gfi1 in D-oDSGCs

Given the difference between *Gfi1* and *Tbx5* conditional mutants, wherein only the downward OKR is affected as opposed to the entire vertical OKR, we sought to determine whether this reflects a greater dependency of the vertical OKR on U-oDSGCs, or rather is reflective of a partial phenotype due to incomplete *Gfi1* removal. To determine D-oDSGC loss in these conditional *Gfi1* mutants, we assessed the ventral MTN innervation in *Chx10^Cre^; Gfi1^F/F^* mice. Unexpectedly, we observed that ventral MTN innervation is completely intact in these mice, suggesting no loss of D-oDSGCs (Figures 4A and 4B). To confirm that D-oDSGCs are indeed present, we labeled Up- and D-oDSGCs by stereotactically injecting CTB-555 retrobeads into the MTN of adult *Chx10^Cre^;Gfi1^F/F^* mice (Figure 4A). We found no difference in the density of MTN-projecting RGCs, nor any difference in the number of RGC couplets[26], in which typically one oDSGC is an U-oDSGC and the other is a D-oDSGC (Figure 4D–4G). To exclude the possibility that there is a loss of D-oDSGCs which is masked by an expansion of U-oDSGCs innervating the ventral MTN, we also examined the number of *Spig1^GFP^* U-oDSGCs in *Chx10^Cre^, Gfi1^F/F^* P10 retinas and found no difference in the number of GFP^+^ RGCs (Figures S4C–S4E). Further, to confirm that these MTN-projecting RGCs indeed maintain D-oDSGC identity, we performed *in situ hybridization* on P6 *Chx10^Cre^; Gfi1^F/F^* retina cryosections. We detected *Gfi1* expression using probes targeting the *Gfi1* locus surrounding the loxP sites, and we found that *Gfi1* and *Fibcd1* are co-expressed in D-oDSGCs in the retinas of conditional *Gfi1* mutants, confirming D-oDSGC identity (Figures S4A and S4B). To investigate whether the discrepancy in D-oDSGC loss between the conditional and global *Gfi1* mutant phenotypes extends to F-mini ONs, we immunostained retinas from *Chx10^Cre^, Gfi1^F/F^* mice for Foxp2.

**Figure 4.**
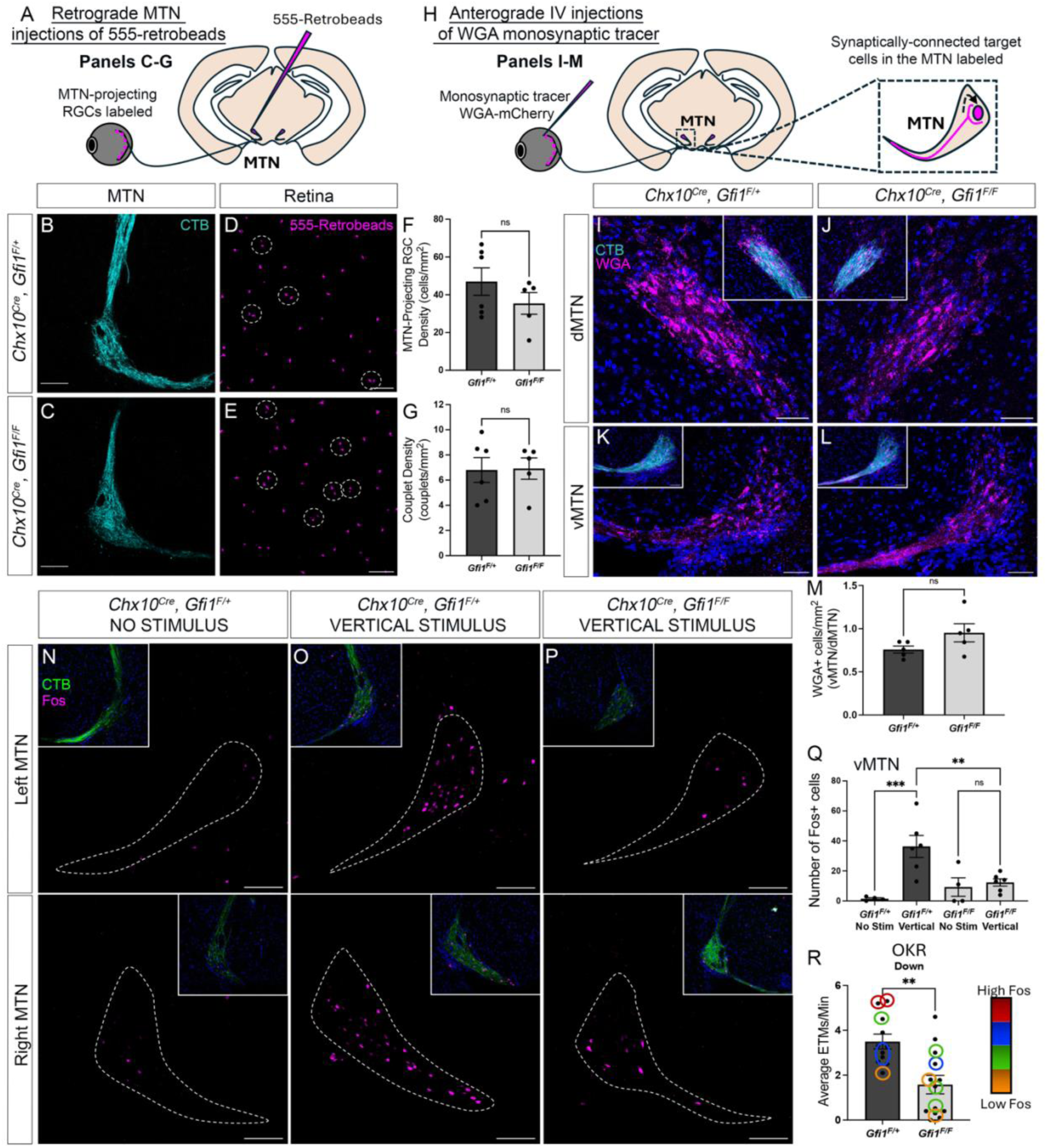
***Gfi1* conditional removal preserves D-oDSGC identity but disrupts function** (**A**) Schematic depicting retrograde injection of 555-retrobeads into the MTN to assess MTN-projecting RGC density (**B, C**) *Chx10^Cre^, Gfi1^F/F^* mice show intact ventral MTN innervation suggesting no loss of D-oDSGCs. (**D-G**) Wholemount retinas of *Chx10^Cre^, Gfi1^F/+^* and *Chx10^Cre^, Gfi1^F/F^* mice following retrograde injection of 555-retrobeads into the MTN confirm no change in MTN-projecting RGC density (**F**) or U- and D-oDSGC couplets (**G**, dashed circles in **D** and **E**). (**H**) Schematic depicting transsynaptic viral labeling in the MTN via intravitral injections of anterograde monosynaptic tracer WGA-mCherry. (**I-M**) Transsynaptic labeling shows no change in WGA^+^ cell density within the MTN upon conditional removal of Gfi1. (**N-Q**) *Chx10^Cre^, Gfi1^F/+^* mice demonstrate strong vertical motion-induced activation of the ventral MTN (outlined in white) estimated from Fos expression (**N, O** and **Q**) unlike *Chx10^Cre^, Gfi1^F/F^* mice which show no ventral MTN activation (**P, Q**). Levels of Fos in the ventral MTN show a strong correlation with strength of the downward OKR response (**R**). Data are presented as mean ± SE. **p < 0.01, ***p < 0.001.

Interestingly, we found a ∼25% reduction in F-RGCs in the retinas of P10 *Chx10^Cre^; Gfi1^F/F^* mice (Figures S4F–S4H), reflecting a loss of F-mini ONs that phenocopies the F-mini ON loss in global *Gfi1* mutants. These distinct phenotypes highlight the importance played by timing of Gfi1 removal, and they suggest that D-oDSGCs are likely generated relatively early in RGC development such that Cre-dependent removal of Gfi1 occurs after the critical window within which their fate is specified, while F-mini ON differentiation occurs later in development. We ruled out the contribution of any allele-specific effects on the different D-oDSGC phenotypes by generating global *Gfi1* null mice from the conditional *Gfi1^Flox^* allele using an embryonically active *Sox2^Cre^* to drive early global Cre-mediated recombination. These *Gfi1^Δ/Δ^* mice showed a complete absence of the ventral MTN tract (Figure S4I–S4K) and a loss of F-mini ONs (Figure S4L), phenocopying the *Gfi1^Cre/Cre^* global null mutant.

Despite the presence of D-oDSGCs, *Chx10^Cre^, Gfi1^F/F^* mice showed significant impairments in the downward OKR response (Figure 3D). We reasoned that the loss of F-mini ONs in these *Gfi1* conditional mice does not contribute to the deficits in downward OKR because: (i) F mini ONs are direction-selective in response to significantly higher speeds than those tested in our OKR assays[30]; and (ii) the AOS is responsible for driving the OKR and F-mini ONs do not project to any AOS nuclei. Thus, the downward OKR impairments seen in *Chx10^Cre^; Gfi1^F/F^* mice strongly suggest a deficit in D-oDSGC function and hint at a post-differentiation role for Gfi1 in D-oDSGC development. Taken together, these results show that our retina-specific knockout of *Gfi1* using the *Chx10^Cre^* line disrupts the specification of F-mini ONs, but preserves D-oDSGC identity, unmasking a Gfi1 post-differentiation function in shaping D-oDSGC development.

### Gfi1 serves a post-differentiation role critical for D-oDSGC function

Generation of D-oDSGCs is normally followed by their integration into retinal circuits. This requires both innervating and assembling synapses with target cells in the ventral MTN, and also elaborating dendrites to establish precise upstream connectivity with starburst amacrine cells and bipolar cells. To determine whether Gfi1 removal following D-oDSGC generation disrupts integration of these DSGCs into the DS circuit, we first examined their synaptic connectivity with target cells in the ventral MTN. We injected *Chx10^Cre^; Gfi1^F/F^* mice intraocularly with an anterograde monosynaptic wheat germ agglutinin (WGA) tagged with mCherry[40] and quantified the number of synaptically-connected target cells in the ventral MTN as compared to the dorsal MTN (Figure 4H). We found no difference in the number of target cells between control and *Chx10^Cre^; Gfi1^F/F^* mice (Figures 4I–4M), suggesting no disruption in the innervation of the ventral MTN nor synaptic connectivity with target cells. These results suggest that any deficits in D-oDSGC computation occur in the retina upstream of the ventral MTN.

To localize the impairment in downward OKR tracking specifically to D-oDSGC functions in the retina, we examined the ability of D-oDSGCs to directly activate the ventral MTN. We assessed the expression of Fos, a neural activity marker, in the ventral MTN of mice in response to continuously-rotating vertical stimuli. Control mice showed a strong upregulation of Fos in the ventral MTN in response to vertical stimuli (Figures 4N, 4O and 4Q), consistent with previous results[26]. However, no such upregulation of Fos was observed in the ventral MTN of *Chx10^Cre^, Gfi1^F/F^* mice (Figures 4P and 4Q), mirroring the disrupted downward OKR in these mice. The levels of Fos in the ventral MTN correlate with the strength of the downward OKR; mice showing greater Fos upregulation also demonstrate stronger downward a OKR (Figure 4R). Collectively, these results localize the OKR behavioral deficit to D-oDSGC function upstream of the ventral MTN.

### A novel role for Gfi1 in regulating D-oDSGC dendritic morphology

To determine how Gfi1 disrupts D-oDSGC function, we assessed the dendritic morphologies of D-oDSGCs. To label the dendrites of vertically-tuned Up- and D-oDSGCs, we stereotactically injected a glycoprotein-deleted rabies virus encoding mCherry into the MTN of P4 *Chx10^Cre^; Gfi1^F/F^* mice (Figure 5A). We used *Spig1^GFP^* to differentiate U-oDSGCs from D-oDSGCs. We assessed dendritic stratification of D-oDSGCs in P7 retina cryosections and found that most control D-oDSGCs stratify primarily in S4, with some extending a small fraction of their dendritic arbor into S2, consistent with previous findings[5] (Figures 5B, 5C, 5F and 5G). Though we found no difference in the stratification of D-oDSGCs in *Chx10^Cre^ Gfi1^F/F^* mice (Figures 5B–5G), many D-oDSGCs showed significantly shorter dendrites (Figure 5H). To better visualize the dendritic arbors of D-oDSGCs, we measured the dendritic field areas of Up- and D-oDSGCs in P7 wholemount retinas (Figures 5I and 5J). While U-oDSGCs showed no difference in dendritic field areas upon pan-retinal loss of Gfi1, D-oDSGCs in *Chx10^Cre^; Gfi1^F/F^* mice had much smaller dendritic fields compared to control D-oDSGCs (Figures 5K–5Q). We also traced the dendrites of D-oDSGCs and found that Gfi1-deficient D-oDSGCs showed significantly reduced total dendritic length and dendritic volume (Figures 5R and 5S), and also less complex dendritic arbors as measured by Sholl analysis (Figure 5T). Finally, we also measured the dendritic field area at P10, after the onset of direction-selectivity, and found a similar reduction in dendritic field area of Gfi1-deficient D-oDSGCs, indicating a persistence of this phenotype following the establishment of directional tuning which occurs ∼P8[41, 42] (Figure S5A). Taken together, these results demonstrate a novel additional function for Gfi1 in regulating the dendritic morphology of D-oDSGCs post-differentiation.

**Figure 5.**
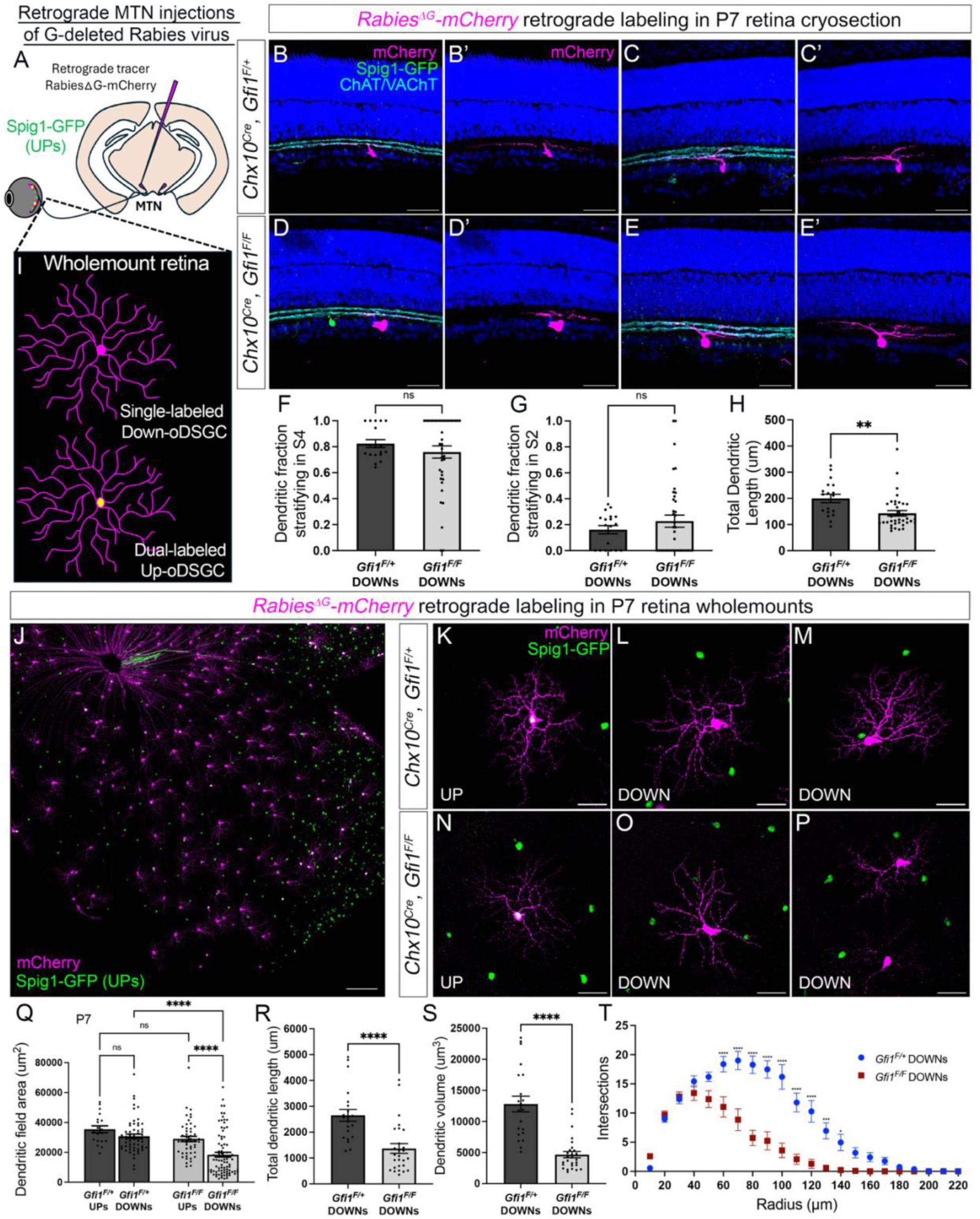
**D-oDSGCs in *Gfi1* conditional mutants show a marked reduction in size and complexity of their dendritic arbors.** (**A**) Schematic of viral retrograde filling of MTN-projecting U-oDSGCs (Spig1^GFP+^) and D-oDSGCs (Spig1^GFP-^). (**B-G**) Spig1^GFP-^D-oDSGCs in *Chx10^Cre^, Gfi1^F/F^* mice show no changes in their dendritic fraction stratifying in either S4 (**F**) or S2 (**G**). S4 and S2 layers are identified by staining for cholinergic SACs that stratify in these layers. (**H**) D-oDSGCs do however display a significant reduction in their total dendritic length. (**I, J**) Schematic (**I**) and P7 wholemount retina (**J**) depicting viral retrograde filling of dual-labeled MTN-projecting U-oDSGCs and single-labeled MTN-projecting D-oDSGCs. (**K-Q**) *Chx10^Cre^, Gfi1^F/F^* mice show similar dendritic field areas of Spig1^GFP+^ U-oDSGCs (**K, N, Q**), but significantly smaller dendritic field areas of Spig1^GFP-^D-oDSGCs (**L, M, O, P, Q**). Many D-oDSGCs display short and poorly elaborated dendrites (**O**). (**R-T**) Dendritic tracing of D-oDSGCs reveals a sharp reduction in total dendritic length (**Q**), dendritic volume (**R**), and complexity (**S**). Data are presented as mean ± SE. *p < 0.05, ***p < 0.001, ****p < 0.0001.

## Discussion

There is a vast diversity in RGC subtypes, yet fundamental questions remain as to how retinal progenitors differentiate into ∼45 distinct RGC subtypes in the murine retina. In this study, we sought to determine the function of the transcription factor Gfi1 in the specification of retinal direction-selective ganglion cells given its strict cell-type specific expression. We found that *Gfi1* is expressed in select RGC populations, including D-oDSGCs and F-mini ONs, both of which exhibit vertical directional tuning. Using a *Gfi1^Cre^* knock-in line to simultaneously disrupt Gfi1 function and to visualize these RGC populations over time, we uncover a critical role for Gfi1 in specifying the fate of both D-oDSGCs (D-oDSGCs) and F-mini ONs. Specifically, we showed that removal of Gfi1 results in improper specification of RGCs and their subsequent loss during the first postnatal week. This developmental trajectory mirrors that seen in inner ear hair cell development[15] where hair cells form initially in *Gfi1* mutants but are morphologically abnormal and subsequently lost in the first two postnatal weeks. Our findings are similar, suggesting that Gfi1 functions in parallel across sensory systems. Additionally, using conditional removal of *Gfi1* we uncovered a novel post-differentiation role for Gfi1 in D-oDSGC development. We generated a retina-specific knockout of *Gfi1* using *Chx10^Cre^* and observed a selective deficit in the downward OKR behavior despite the presence of D-oDSGCs. We localized this deficit to D-oDSGC function by observing reduced Fos activation in the ventral, but not the dorsal, MTN. Finally, we showed that D-oDSGCs in these conditional *Gfi1* mutants exhibit altered dendritic morphologies, with significantly smaller and less complex dendritic arbors. Altogether, these results describe a novel dual-function for Gfi1 in regulating the specification and dendritic development of D-oDSGCs, two functions that are segregated across developmental time (Figure 6).

**Figure 6.**
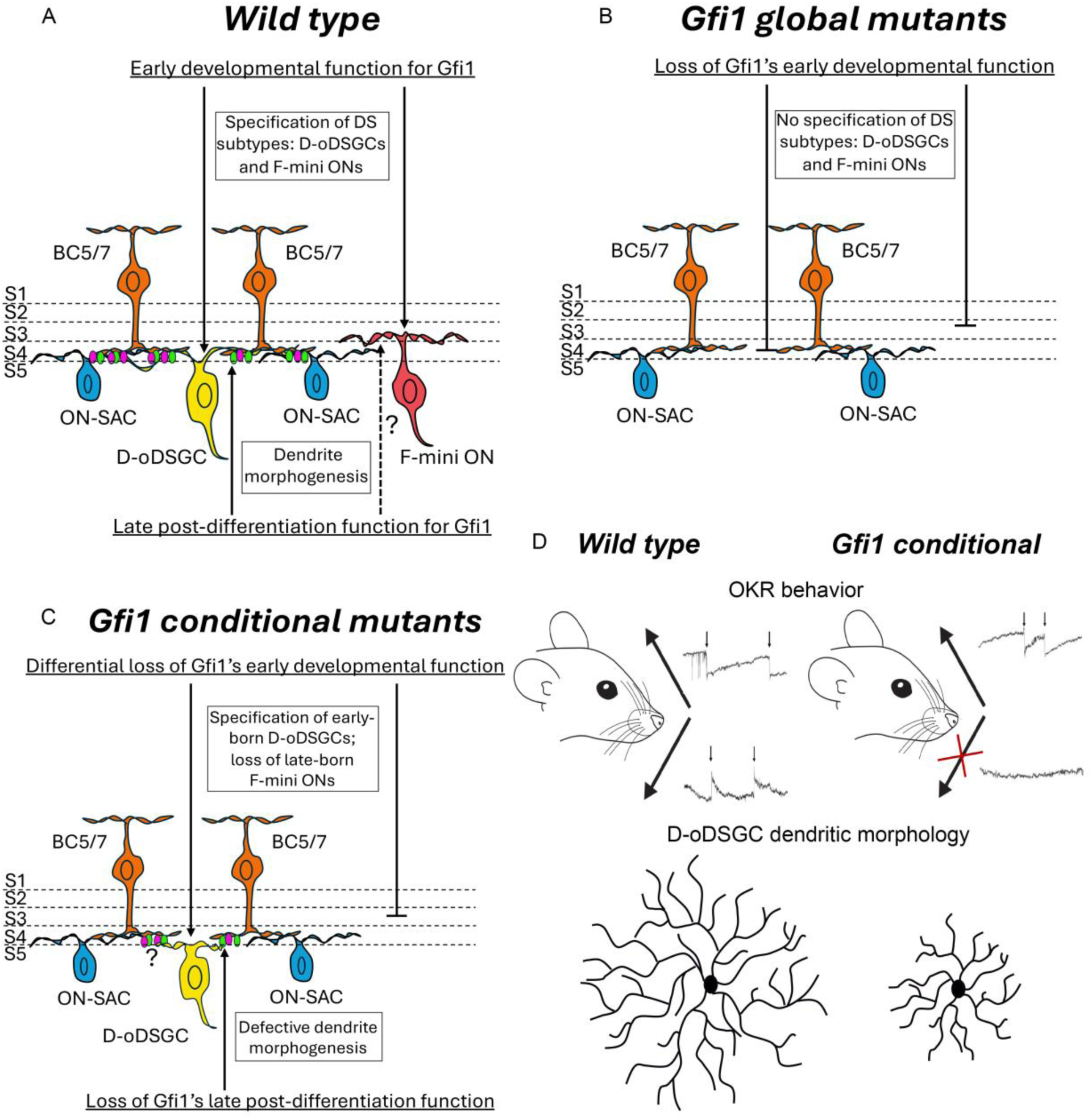
**Summary of Gfi1 functions in retinal direction-selective circuits** (**A**) Schematic representation of the dual functions of Gfi1 in (i) early specification of direction-selective D-oDSGCs and F-mini ONs, and (ii) late dendritic development of D-oDSGCs and possibly, F-mini ONs. (**B**) Loss of Gfi1’s early function in *Gfi1* global mutants results in a specific loss of direction-selective D-oDSGCs and F-mini ONs. (**C**) Conditional *Gfi1* mutants show a differential loss of Gfi1’s early function where late-born F-mini ONs are lost but early-born D-oDSGCs persist, revealing a late post-differentiation function of Gfi1 in regulating D-oDSGCs dendritic development and possibly their upstream synaptic connectivity. (**D**) Summary of the phenotypes seen in conditional *Gfi1* mutants including loss of downward motion detection in OKR behavior and smaller and less branched D-oDSGC dendritic arbors.

The post-differentiation function for Gfi1was revealed by the retina-specific knockout of *Gfi1* that we generated using *Chx10^Cre^* to avoid confounding effects on OKR behavior resulting from VOR deficits in global *Gfi1* mutants. *Chx10^Cre^* is a well-characterized line that expresses Cre in retinal progenitor cells beginning at ∼E10.5[36]. With this line, we found a differential effect of conditional *Gfi1* removal on D-oDSGCs and F-mini ONs, wherein the former population is preserved but dysfunctional while the latter population is lost. We attribute this differential effect to timing of their specification, with D-oDSGCs likely differentiating before F-mini ONs. Indeed, *Spig1^GFP^* U-oDSGCs are already formed by E13.5[22], and previous reports show differences in birth timing for certain RGC subtypes[43]. However, a more comprehensive characterization of birth timings of all RGC subtypes will provide insight into this issue. Further, we find no evidence that mosaicism of *Chx10^Cre^* underlies these differences since we observe >95% recombination efficiency using the *Ai14* reporter. We also find no contribution from allele-specific effects since global mutants generated from the *Gfi1* conditional allele phenocopy global *Gfi1^Cre/Cre^* mutants. Collectively, these data support the notion that timing of *Gfi1* removal underlies the differential phenotypes of D-oDSGCs and F-mini ONs in our conditional mutants.

D-oDSGCs in the *Gfi1* conditional mutants show altered dendritic morphologies and a loss of downward motion encoding, as evidenced by a lack of ventral MTN activation. What is the mechanism underlying this loss of downward motion detection? One possibility is that altered dendritic morphology reflects disruptions in the precise upstream connectivity with starburst amacrine cells and bipolar cells, influencing the proper establishment of direction-selectivity in D-oDSGCs. Alternatively, smaller dendritic arbors might prevent the active dendritic integration underlying computation of direction-selectivity[44]. The presence of these dysfunctional D-oDSGCs in *Gfi1* conditional retinas presents the unique opportunity to identify downstream effectors of Gfi1 that underlie directional tuning, molecules which to date have remained elusive.

We find a conserved role for Gfi1 in the differentiation of two direction-selective RGC subtypes – D-oDSGCs and F-mini ONs. Though we have not directly assessed whether the third Gfi1-expressing subtype, W3B[29], is lost in global *Gfi1* mutants, the persistence of ∼50% of Gfi1-expressing RGCs suggests that they are unaffected by the loss of *Gfi1*. If so, these results suggest a shared function for Gfi1 that is specific to D-oDSGCs and F-mini ONs. Unlike W3Bs, both D-oDSGCs and F-mini ONs are direction-selective, primarily tuned to vertical motion, and it is possible that Gfi1 serves to regulate a conserved pathway for the establishment of downward direction-selectivity. Notably, despite F-mini ONs accounting for ∼40% of F-RGCs, *Gfi1* is expressed in only 25% of F-RGCs, showing that *Gfi1*-expression is limited to only a subset of F-mini ONs. This is especially interesting since F-mini ONs exist as two populations themselves: those in the dorsal retina preferring ventral motion, and those in the ventral retina preferring dorsal motion[30]. Future work will explore whether *Gfi1* expression is restricted to downward-preferring F-mini ONs and whether Gfi1 also has a post-differentiation function in regulating the dendritic morphologies and directional tuning of downward-preferring F-mini ONs.

In both *Drosophila* R8 photoreceptors and mouse inner ear hair cells, Senseless/Gfi1 is necessary and sufficient to drive their respective fates[15, 18, 19, 45, 46]. In *Drosophila* photoreceptors, Senseless functions with Atonal to direct R8 fate[19], while Gfi1 together with Atoh1 and Pou4f3 can reprogram cells to adopt the hair cell fate[45, 46]. In the mouse retina, Atoh7 and Pou4f2 play important roles in the specification of RGCs[47–51], though neither is specific for the differentiation of any one RGC subtype. It will be interesting to assess whether coupling the expression of *Gfi1* with these other transcription factors allows for the specification of D-oDSGCs and F-mini ONs, selectively.

Collectively, our findings describe novel roles for the transcription factor Gfi1 in specifying down DS fates of D-oDSGCs and F-mini ONs. Additionally, we describe a novel late function for Gfi1 in regulating the dendritic development and function of D-oDSGCs. The implication that Gfi1 is necessary for the maturation of down direction-selective RGCs specifically highlights its importance as a candidate for revealing fundamental mechanisms underlying direction-selective specification and tuning.

## Data availability Lead Contact

Further information and requests for resources and reagents should be directed to and will be fulfilled by the lead contact, Alex L. Kolodkin.

## Acknowledgments

We are grateful to Masaharu Noda, Ulrich Mueller and Seth Blackshaw for generously providing *Spig1^GFP^, Gfi1^Cre^*, *Gfi1^flox^*, *Gfi1^CreERT2^* and *Chx10^Cre^* mice, respectively. We thank all members of the Kolodkin laboratory for comments and helpful discussions.

This work was supported by R01 EY032095 (ALK).

## Methods

### Experimental model and subject details

All animal experiments were approved by the Institutional Animal Care and Use Committee (IACUC) at The Johns Hopkins University School of Medicine. The day of birth was designated as postnatal day 0 (P0). Mice of both sexes were used. *Spig1-GFP* mice were a gift from M. Noda and were described previously[25, 26]. *Gfi1^Cre^*, *Gfi1^flox^* and *Gfi1^CreERT2^* mice were gifts from U. Mueller and were described previously[23, 34, 35]. *Chx10^Cre^* mice were a gift from S. Blackshaw and were described previously[36]. *Sox2^Cre^* mice were obtained from Jackson Laboratories. All mice were maintained on a C57BL/6J background. Animals were housed in a 12-hour light-dark cycle. Behavioral tests were performed at consistent hours during the light cycle.

### Intraocular CTB and AAV injections

Mice were anaesthetized with 2% isoflurane. A hole was made at the corneal limbus with a 30 G needle and 1 ul of CTB or AAV was injected intravitreally using a Hamilton syringe. For co-injections, 0.75 ul of CTB with 0.75 ul of AAV were co-injected.

The following viruses were used in this study: AAV9 EF1a-BbChT (>1E+13; Addgene #45186-AAV9) and AAV2 CAG-mWGA-mCherry (2E+12; Virovek, Inc. Cat # 117BUCSF89).

### Stereotactic surgery and MTN injections

P4 *Spig1-GFP* mice were anesthetized using 2% isoflurane and placed into a stereotactic apparatus. Rabies virus (SAD-B19-RVdG-tdTomato (3.50E+10; UC Irvine Vector Core, Cat # RVdG-5) was injected into the medial terminal nucleus (100 nl at 10 nl/s) using a Hamilton Neuros syringe coupled to a computerized microsyringe pump controller (World Precision Instruments). Five minutes elapsed between the injection and removal of the needle to allow diffusion of CTB around the injection site. The MTN coordinates at P4 are as follows: anterior/posterior -1.13 mm, medial/lateral ±1.0 mm, dorsal/ventral -5.01 mm, needle tilted 50° anteriorly (coordinates are relative to the intersection of the superior sagittal sinus and the inferior cerebral vein). Following injection, five minutes were allowed to elapse before removing the syringe to allow for diffusion to occur. Labeled neurons were traced using the Neurolucida 360 software from MBF Bioscience.

Adult mice were anesthetized using 2% isoflurane and placed into a stereotaxic apparatus. Lumafluor-555 retrobeads were injected into the medial terminal nucleus (600 nL at 10 nL/s) using a Hamilton Neuros syringe via an automatic microsyringe controller (World Precision Instruments). The MTN was injected at the following coordinates relative to bregma: anterior/posterior 0 mm, medial/lateral ± 0.85 mm, dorsal/ventral -5.4mm with a needle tilt of 30° anteriorly. Needle was brought to lowest dorsal/ventral depth, raised 0.2mm, and injected. Following injection, five minutes were allowed to elapse before removing the syringe to allow for diffusion to occur.

### Immunohistochemistry

For whole-mount retina staining, mice were transcardially perfused with phosphate buffered saline (PBS) followed by 4% paraformaldehyde (PFA). Enucleated eyeballs were fixed in 4% PFA for 1 hour at 4°C, then washed 3 times with PBS to remove residual PFA. Retinas were dissected and incubated for 2 days at 4°C with primary antibodies in PBS containing 0.4% Triton X-100, 5% donkey (or goat) serum, and 0.05% sodium azide. Retinas were washed 4 times for 1 hour at room temperature with PBS and 0.1% Triton X-100, then incubated for overnight at 4°C with secondary antibodies in PBS containing 0.4% Triton X-100 and 5% donkey (or goat) serum. Retinas were washed 4 times for 1 hour at room temperature with PBS and 0.4% Triton X-100, then mounted and imaged with a Zeiss LSM 700 confocal microscope.

For cross-sectional retina staining, mice were transcardially perfused with PBS followed by 4% PFA. Enucleated eyeballs were fixed in 4% PFA for 1 hour at 4°C, then washed 3 times with PBS to remove residual PFA. A hole was made in the cornea and the eyeballs were cryopreserved in PBS containing 30% (w/v) sucrose overnight at 4°C. Eyeballs were frozen in Neg-50 frozen section medium (Richard-Allen Scientific, Kalamazoo, MI) and sliced using a cryostat at a thickness of 20 um. Retinal sections were blocked in PBS containing 5% donkey (or goat) serum and 0.4% Triton X-100 at room temperature for 1 hour, then incubated with primary antibodies in PBS containing 5% donkey (or goat) serum and 0.4% Triton X-100 overnight at 4°C. Sections were washed 6 times for 5 minutes at room temperature with PBS and 0.1% Triton X-100, then incubated with secondary antibodies in PBS containing 5% donkey (or goat) serum and 0.4% Triton X-100 at room temperature for 1 hour. Sections were washed 6 times for 5 minutes at room temperature with PBS and 0.1% Triton X-100, then mounted for imaging.

For cross-sectional brain staining, mice were transcardially perfused with PBS followed by 4% PFA. Brains were fixed in 4% PFA overnight at 4°C, then washed 3 times with PBS to remove residual PFA. Brains were embedded in PBS containing 4% (w/v) agarose and sliced using a vibratome at a thickness of 200 um. Brain sections were blocked in PBS containing 2% bovine serum albumin (BSA) and 0.1% Triton X-100 and 0.1% Tween-20 at room temperature for 1 hour, then incubated with primary antibodies in PBS containing 2% BSA and 0.1% Tween-20 for overnight at 4°C. Sections were washed 4 times for 1 hour at room temperature with PBS and 0.1% Tween-20, then incubated with secondary antibodies in PBS containing 2% BSA and 0.1% Tween-20 for overnight at 4°C. Sections were washed 4 times for 1 hour at room temperature with PBS and 0.1% Tween-20, then mounted for imaging.

Primary antibodies used in this study include: Rabbit anti-dsRed (Living Colors, 1:1000), Chick anti-GFP (AVES, 1:1000), Guinea Pig anti-Foxp2 (Synaptic Systems, 1:500), Rabbit anti-Rbpms (Proteintech, 1:500), Guinea Pig anti-Rbpms (ThermoFisher, 1:500), Mouse anti-Brn3c (Santa Cruz, 1:100), Rabbit anti-Foxp1 (Proteintech, 1:500), Rabbit anti-IBA1 (Wako Laboratory Chemicals, 1:500), Goat anti-ChAT (Abcam, 1:250), Goat anti-VAChT (Millipore, 1:250) and Rabbit anti-cFos (Cell Signaling Technology, 1:500).

### *In situ* hybridization

Fluorescent *in situ* hybridization was performed on fresh frozen retinas from P4 and P6 mice. Animals were anesthetized using 2% isoflurane and decapitated. Enucleated eyeballs were embedded in Neg-50 frozen section medium (Richard-Allen Scientific, Kalamazoo, MI), frozen on dry ice, and sliced using a cryostat at a thickness of 14 um. *In situ* hybridization was performed using the RNAscope Fluorescent Multiplex system v1 (Advanced Cell Diagnostics, Newark, CA) or Hybridization Chain Reaction RNA-FISH Technology from Molecular Instruments according to the manufacturer’s instructions. Probes for *Gfi1*, *Tbx5*, *Rbpms,* and *Fibcd1* were obtained from the manufacturer (Advanced Cell Diagnostics, Newark, CA). Dotted lines were drawn around the collection of mRNA puncta for the gene of interest. In our experience, the mRNA puncta were often located outside of the DAPI^+^ nucleus, but were grouped in a circular shape that presumably corresponds to the cell soma. Punctate dots within cells indicated positive staining.

### Headpost implantation surgery

Adult mice (≥6 weeks of age) were anesthetized using isoflurane. Four 1.00UNM × 0.120″ stainless steel screws were placed into the skull on both sides of the sagittal suture between the coronal and lambdoidal sutures. The screws were incorporated into a pedestal of dental cement (Ortho-Jet, Lang Dental; Wheeling, Illinois, USA) and an acrylic headpost, composed of two M1.4 hex nuts embedded in dental cement (fabricated in-house), was placed on top of the pedestal. Mice were allowed to recover for ≥7 days before behavioral testing.

### Optokinetic reflex recording

Mice with headposts were restrained in an animal holder (fabricated in-house) and were placed inside a square box, each side of which consisted of a computer monitor (12 in length × 20 in width) displaying a black-and-white checkerboard pattern (stripe width, 5° of visual angle). Mice were positioned so that their two eyes saw different pairs of monitors, allowing presentation of visual rotational stimuli around the anterior-posterior axis. A fixed infrared video camera inside the box recorded eye movements. Linear displacement of the pupil in the 2D video image (in mm) was transformed into rotational displacement (in degrees) using the video-oculography method described previously[22]. Two types of visual stimuli were used to elicit the optokinetic reflex: sinusoidal stimuli (temporal frequency, 0.1 or 0.2 Hz; sinusoid amplitude, 5° of visual angle; 10 cycles per trial) and continuously rotating stimuli (rotation speed, 5° of visual angle/s; each stimulus cycle consisted of 30 s of rotating checkerboard, followed by 30 seconds of grey screen; 10 cycles per trial). Stimuli were presented in both vertical and horizontal directions. Eye movement recording and stimulus presentation programs were built in-house using MATLAB R2012a v7.14.0.739 (MathWorks, Natick, MA) and LabVIEW v12.0f3 (National Instruments, Austin, TX) software. Data were analyzed using Igor Pro v6.37 (WaveMetrics, Portland, OR) as described previously[22].

### Quantification and statistical analysis

Graphs were generated using the ggplot2 package in R v3.5.1 (The R Foundation for Statistical Computing, Auckland, New Zealand) and GraphPad Prism 9.5.0. Significance was determined with either a *t*-test or a one-way ANOVA with a post-hoc Dunnet’s test. Significance was defined as p < 0.05.

## References

1. Kodama, T. and S. du Lac, Adaptive Acceleration of Visually Evoked Smooth Eye Movements in Mice. J Neurosci, 2016. 36(25): p. 6836–49.

2. Harris, S.C. and F.A. Dunn, Asymmetric retinal direction tuning predicts optokinetic eye movements across stimulus conditions. Elife, 2023. 12.

3. Vaney, D.I., B. Sivyer, and W.R. Taylor, Direction selectivity in the retina: symmetry and asymmetry in structure and function. Nat Rev Neurosci, 2012. 13(3): p. 194–208.

4. Hamilton, N.R., A.J. Scasny, and A.L. Kolodkin, Development of the vertebrate retinal direction-selective circuit. Dev Biol, 2021. 477: p. 273–283.

5. Dhande, O.S., et al., Genetic dissection of retinal inputs to brainstem nuclei controlling image stabilization. J Neurosci, 2013. 33(45): p. 17797–813.

6. Sabbah, S., et al., A retinal code for motion along the gravitational and body axes. Nature, 2017. 546(7659): p. 492–497.

7. Simpson, J.I., The accessory optic system. Annu Rev Neurosci, 1984. 7: p. 13–41.

8. Giolli, R.A., R.H. Blanks, and F. Lui, The accessory optic system: basic organization with an update on connectivity, neurochemistry, and function. Prog Brain Res, 2006. 151: p. 407–40.

9. Gilks, C.B., et al., Progression of interleukin-2 (IL-2)-dependent rat T cell lymphoma lines to IL-2-independent growth following activation of a gene (Gfi-1) encoding a novel zinc finger protein. Mol Cell Biol, 1993. 13(3): p. 1759–68.

10. Karsunky, H., et al., Inflammatory reactions and severe neutropenia in mice lacking the transcriptional repressor Gfi1. Nat Genet, 2002. 30(3): p. 295–300.

11. Yücel, R., et al., The transcriptional repressor Gfi1 affects development of early, uncommitted c-Kit+ T cell progenitors and CD4/CD8 lineage decision in the thymus. J Exp Med, 2003. 197(7): p. 831–44.

12. Hock, H., et al., Intrinsic requirement for zinc finger transcription factor Gfi-1 in neutrophil differentiation. Immunity, 2003. 18(1): p. 109–20.

13. Hock, H., et al., Gfi-1 restricts proliferation and preserves functional integrity of haematopoietic stem cells. Nature, 2004. 431(7011): p. 1002–7.

14. Zeng, H., et al., Transcription factor Gfi1 regulates self-renewal and engraftment of hematopoietic stem cells. Embo j, 2004. 23(20): p. 4116–25.

15. Wallis, D., et al., The zinc finger transcription factor Gfi1, implicated in lymphomagenesis, is required for inner ear hair cell differentiation and survival. Development, 2003. 130(1): p. 221–32.

16. Bjerknes, M. and H. Cheng, Cell Lineage metastability in Gfi1-deficient mouse intestinal epithelium. Dev Biol, 2010. 345(1): p. 49–63.

17. Shroyer, N.F., et al., Gfi1 functions downstream of Math1 to control intestinal secretory cell subtype allocation and differentiation. Genes Dev, 2005. 19(20): p. 2412–7.

18. Nolo, R., L.A. Abbott, and H.J. Bellen, Senseless, a Zn finger transcription factor, is necessary and sufficient for sensory organ development in Drosophila. Cell, 2000. 102(3): p. 349–62.

19. Xie, B., et al., Senseless functions as a molecular switch for color photoreceptor differentiation in Drosophila. Development, 2007. 134(23): p. 4243–53.

20. Jen, H.I., et al., GFI1 regulates hair cell differentiation by acting as an off-DNA transcriptional co-activator of ATOH1, and a DNA-binding repressor. Sci Rep, 2022. 12(1): p. 7793.

21. Fraszczak, J., et al., Reduced expression but not deficiency of GFI1 causes a fatal myeloproliferative disease in mice. Leukemia, 2019. 33(1): p. 110–121.

22. Al-Khindi, T., et al., The transcription factor Tbx5 regulates direction-selective retinal ganglion cell development and image stabilization. Curr Biol, 2022. 32(19): p. 4286–4298.e5.

23. Yang, H., et al., Gfi1-Cre knock-in mouse line: A tool for inner ear hair cell-specific gene deletion. Genesis, 2010. 48(6): p. 400–6.

24. Wässle, H. and H.J. Riemann, The mosaic of nerve cells in the mammalian retina. Proc R Soc Lond B Biol Sci, 1978. 200(1141): p. 441–61.

25. Yonehara, K., et al., Expression of SPIG1 reveals development of a retinal ganglion cell subtype projecting to the medial terminal nucleus in the mouse. PLoS One, 2008. 3(2): p. e1533.

26. Yonehara, K., et al., Identification of retinal ganglion cells and their projections involved in central transmission of information about upward and downward image motion. PLoS One, 2009. 4(1): p. e4320.

27. Rheaume, B.A., et al., Single cell transcriptome profiling of retinal ganglion cells identifies cellular subtypes. Nat Commun, 2018. 9(1): p. 2759.

28. Tran, N.M., et al., Single-Cell Profiles of Retinal Ganglion Cells Differing in Resilience to Injury Reveal Neuroprotective Genes. Neuron, 2019. 104(6): p. 1039–1055.e12.

29. Zhang, Y., et al., The most numerous ganglion cell type of the mouse retina is a selective feature detector. Proc Natl Acad Sci U S A, 2012. 109(36): p. E2391–8.

30. Rousso, D.L., et al., Two Pairs of ON and OFF Retinal Ganglion Cells Are Defined by Intersectional Patterns of Transcription Factor Expression. Cell Rep, 2016. 15(9): p. 1930–44.

31. Matern, M., et al., Gfi1(Cre) mice have early onset progressive hearing loss and induce recombination in numerous inner ear non-hair cells. Sci Rep, 2017. 7: p. 42079.

32. Shekhar, K., et al., Comprehensive Classification of Retinal Bipolar Neurons by Single-Cell Transcriptomics. Cell, 2016. 166(5): p. 1308–1323.e30.

33. Yan, W., et al., Mouse Retinal Cell Atlas: Molecular Identification of over Sixty Amacrine Cell Types. J Neurosci, 2020. 40(27): p. 5177–5195.

34. Tang, Q., et al., Gfi1-GCE inducible Cre line for hair cell-specific gene manipulation in mouse inner ear. Genesis, 2019. 57(9): p. e23304.

35. Zhu, J., et al., Gfi-1 plays an important role in IL-2-mediated Th2 cell expansion. Proc Natl Acad Sci U S A, 2006. 103(48): p. 18214–9.

36. Rowan, S. and C.L. Cepko, Genetic analysis of the homeodomain transcription factor Chx10 in the retina using a novel multifunctional BAC transgenic mouse reporter. Dev Biol, 2004. 271(2): p. 388–402.

37. Clement, G. and A. Berthoz, Cross-coupling between horizontal and vertical eye movements during optokinetic nystagmus and optokinetic afternystagmus elicited in microgravity. Acta Otolaryngol, 1990. 109(3-4): p. 179–87.

38. Garbutt, S., et al., Disorders of vertical optokinetic nystagmus in patients with ocular misalignment. Vision Res, 2003. 43(3): p. 347–57.

39. Economides, J.R., et al., Vertical Optokinetic Stimulation Induces Diagonal Eye Movements in Patients with Idiopathic Infantile Nystagmus. Invest Ophthalmol Vis Sci, 2020. 61(6): p. 14.

40. Tsai, N.Y., et al., Trans-Seq maps a selective mammalian retinotectal synapse instructed by Nephronectin. Nat Neurosci, 2022. 25(5): p. 659–674.

41. Yonehara, K., et al., Spatially asymmetric reorganization of inhibition establishes a motion-sensitive circuit. Nature, 2011. 469(7330): p. 407–10.

42. Wei, W., et al., Development of asymmetric inhibition underlying direction selectivity in the retina. Nature, 2011. 469(7330): p. 402–6.

43. Osterhout, J.A., et al., Birthdate and outgrowth timing predict cellular mechanisms of axon target matching in the developing visual pathway. Cell Rep, 2014. 8(4): p. 1006–17.

44. Sivyer, B. and S.R. Williams, Direction selectivity is computed by active dendritic integration in retinal ganglion cells. Nat Neurosci, 2013. 16(12): p. 1848–56.

45. Iyer, A.A., et al., Cellular reprogramming with ATOH1, GFI1, and POU4F3 implicate epigenetic changes and cell-cell signaling as obstacles to hair cell regeneration in mature mammals. Elife, 2022. 11.

46. McGovern, M.M., et al., Expression of Atoh1, Gfi1, and Pou4f3 in the mature cochlea reprograms nonsensory cells into hair cells. Proc Natl Acad Sci U S A, 2024. 121(5): p. e2304680121.

47. Brown, N.L., et al., Math5 is required for retinal ganglion cell and optic nerve formation. Development, 2001. 128(13): p. 2497–508.

48. Wang, S.W., et al., Requirement for math5 in the development of retinal ganglion cells. Genes Dev, 2001. 15(1): p. 24–9.

49. Qiu, F., H. Jiang, and M. Xiang, A comprehensive negative regulatory program controlled by Brn3b to ensure ganglion cell specification from multipotential retinal precursors. J Neurosci, 2008. 28(13): p. 3392–403.

50. Badea, T.C., et al., Distinct roles of transcription factors brn3a and brn3b in controlling the development, morphology, and function of retinal ganglion cells. Neuron, 2009. 61(6): p. 852–64.

51. Brodie-Kommit, J., et al., Atoh7-independent specification of retinal ganglion cell identity. Sci Adv, 2021. 7(11).

