## Supplemental Information for "Temporally-segregated dual functions for Gfi1 in the development of retinal direction-selectivity"

**Figure S1**

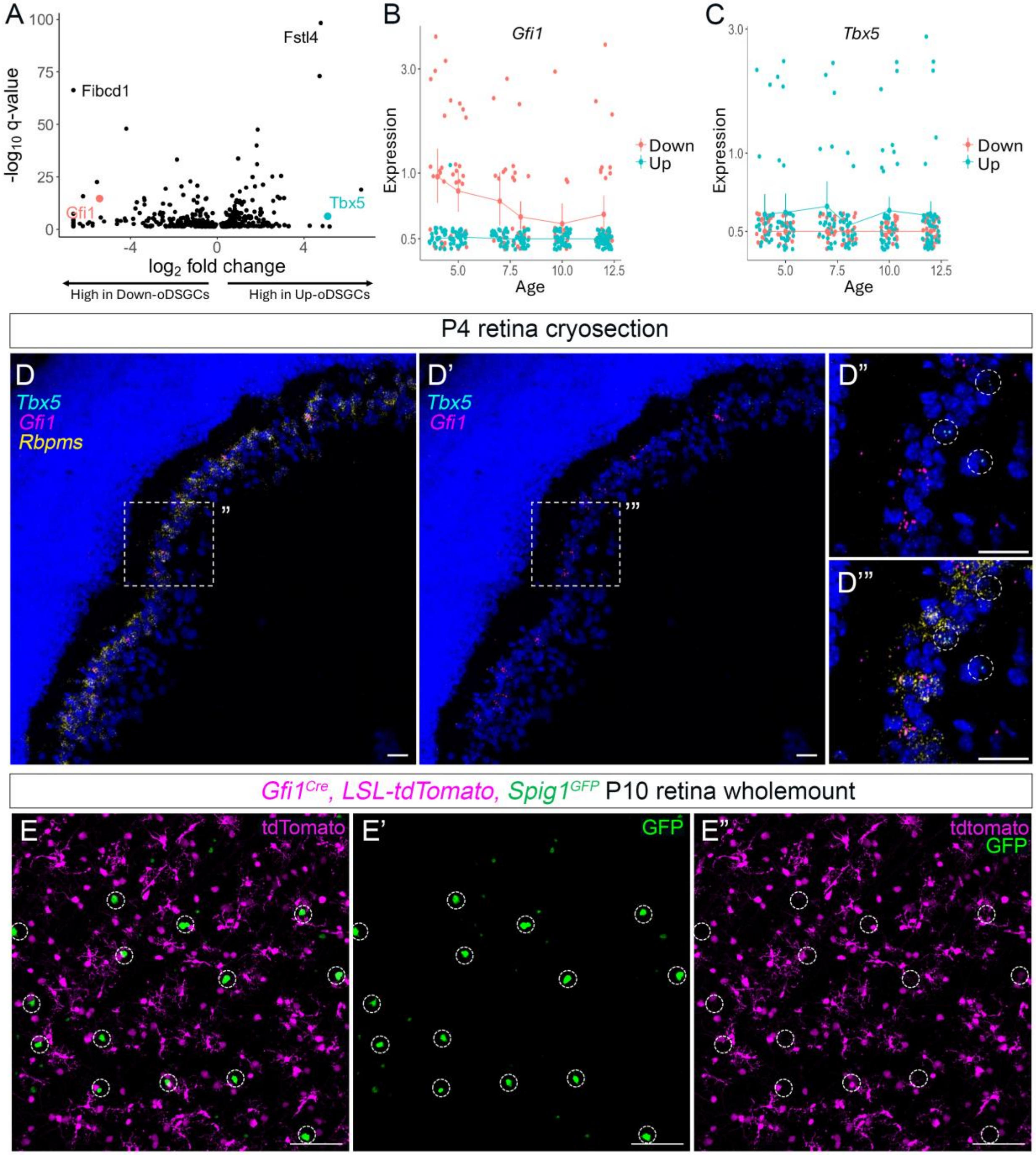

A
Fstl4
Fibcd1
Gfi1
Tbx5
-log10 q-value
log2 fold change
High in Down-oDSGCs
High in Up-oDSGCs
B
Gfi1
Expression
Age
Down
Up
C
Tbx5
Expression
Age
Down
Up
P4 retina cryosection
D
Tbx5
Gfi1
Rbpms
D'
D''
D'''
Gfi1Cre, LSL-tdTomato, Spig1GFP P10 retina wholemount
E
tdTomato
E'
GFP
E''
tdtomato
GFP

**Figure S1. *Gfi1* is selectively expressed in Down- and not U-oDSGCs**

(**A**) Volcano plot showing q values and effect sizes for genes that are significantly differentially expressed between Up- and D-oDSGCs (all genes shown have $q < 0.05$). Fstl4 (Spig1) and Tbx5 were differentially expressed in U-oDSGCs while Fibcd1 and Gfi1 were differentially expressed in D-oDSGCs. (**B, C**) *Gfi1* and *Tbx5* expression at different postnatal ages in Down- and U-oDSGCs, respectively. *Gfi1* expression declines during the end of the first postnatal week. Note the low yet comparable levels of expression between *Gfi1* and *Tbx5.* (**D, D'**) *In situ* hybridization in P4 retina cryosection shows *Gfi1* expression exclusively in *Rbpms*$^{+}$ RGCs, and absent in *Tbx*$^{+}$ U-oDSGCs. (**D", D'"**) Detailed views of the nuclei indicated in D and D'. (**E**) Wholemount P10 *Gfi1*$^{Cre}$*, LSL-tdTomato, Spig1*$^{GFP}$ retina shows mututally exclusive expression of tdTomato and GFP confirming absence of Gfi1 expression in GFP$^{+}$ U-oDSGCs.

**Figure S2**

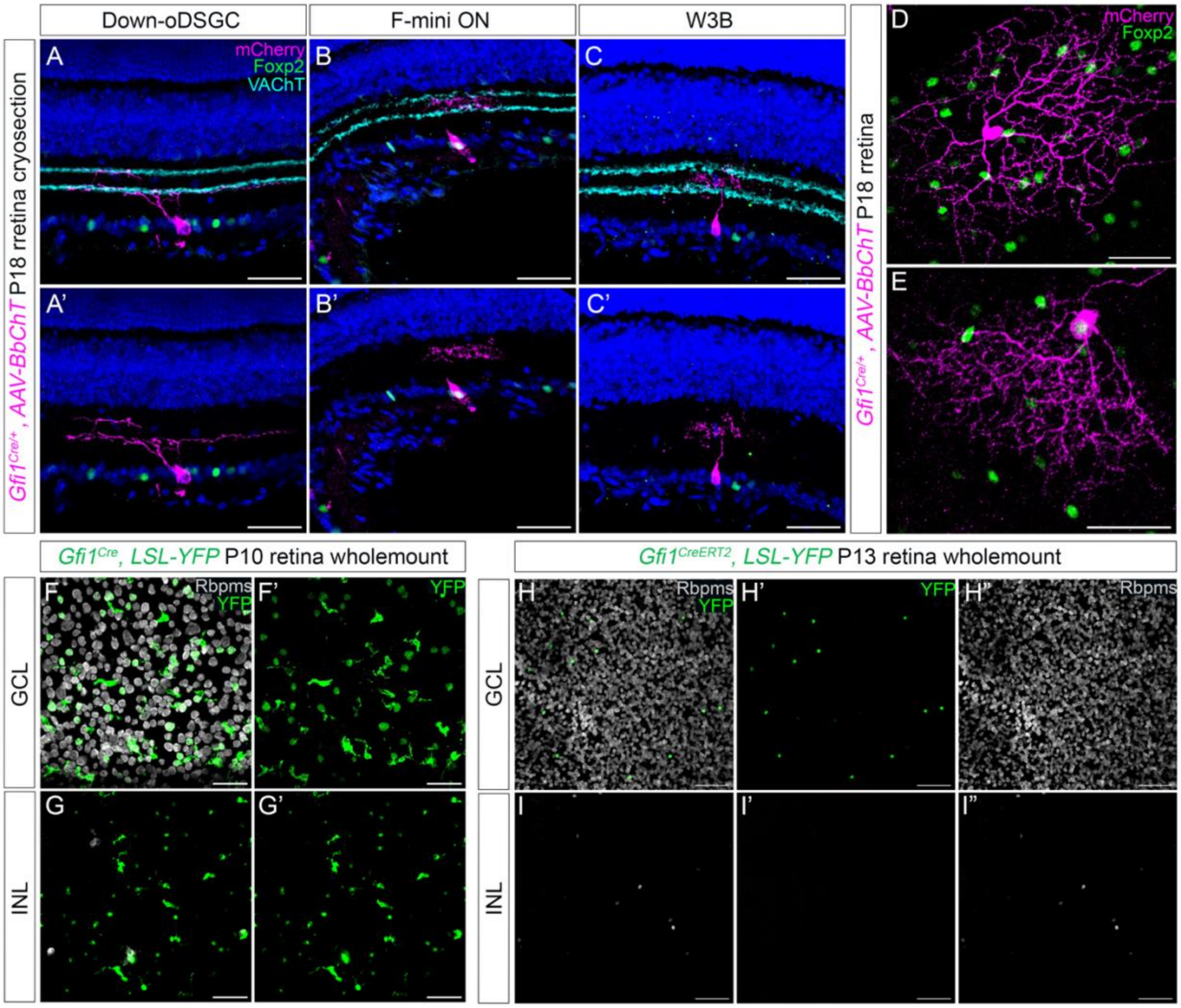

Down-oDSGC
F-mini ON
W3B
mCherry
Foxp2
VAChT
A
B
C
A'
B'
C'
D
E
Gfi1Cre/+, AAV-BbChT P18 rretina cryosection
Gfi1Cre/+, AAV-BbChT P18 rretina
Gfi1Cre, LSL-YFP P10 retina wholemount
Gfi1CreERT2, LSL-YFP P13 retina wholemount
Rbpms
YFP
F
F'
G
G'
H
H'
H"
I
I'
I"
GCL
INL

**Figure S2. *Gfi1* is expressed in direction-selective F-mini ON RGCs**

(**A-C**) Intraocular injection of *AAV9-Brainbow-mCherry* (*AAV-BbChT*) into $Gfi1^{Cre}$ mice reveals three morphologically distinct RGC subtypes corresponding to $Foxp2^-$ D-oDSGCs co-stratifying with SACs in S4 (**A, A'**), $Foxp2^+$ F-mini ONs with bistratified morphology in S3 (**B, B'**), and $Foxp2^-$ W3Bs with dense dendritic arborization in S3 (**C, C'**). (**D, E**) *En face* view of the highly asymmetric dendrites of F-mini ONs (**E**) in contrast to D-oDSGCs (**D**). (**F, G**) Wholemount P10 $Gfi1^{Cre}$*, LSL-YFP* retina shows aberrant expression of *Gfi1* in some $Rbpms^-$ cells in the INL arising after P6. (**H, I**) Wholemount P13 $Gfi1^{CreERT2}$*, LSL-YFP* retina following tamoxifen injections from P5 to P13 shows *Gfi1* expression restricted to $Rbpms^+$ RGCs and absent in the INL, confirming that the aberrant INL expression is due to $Gfi1^{Cre}$ haploinsufficiency.

**Figure S3**

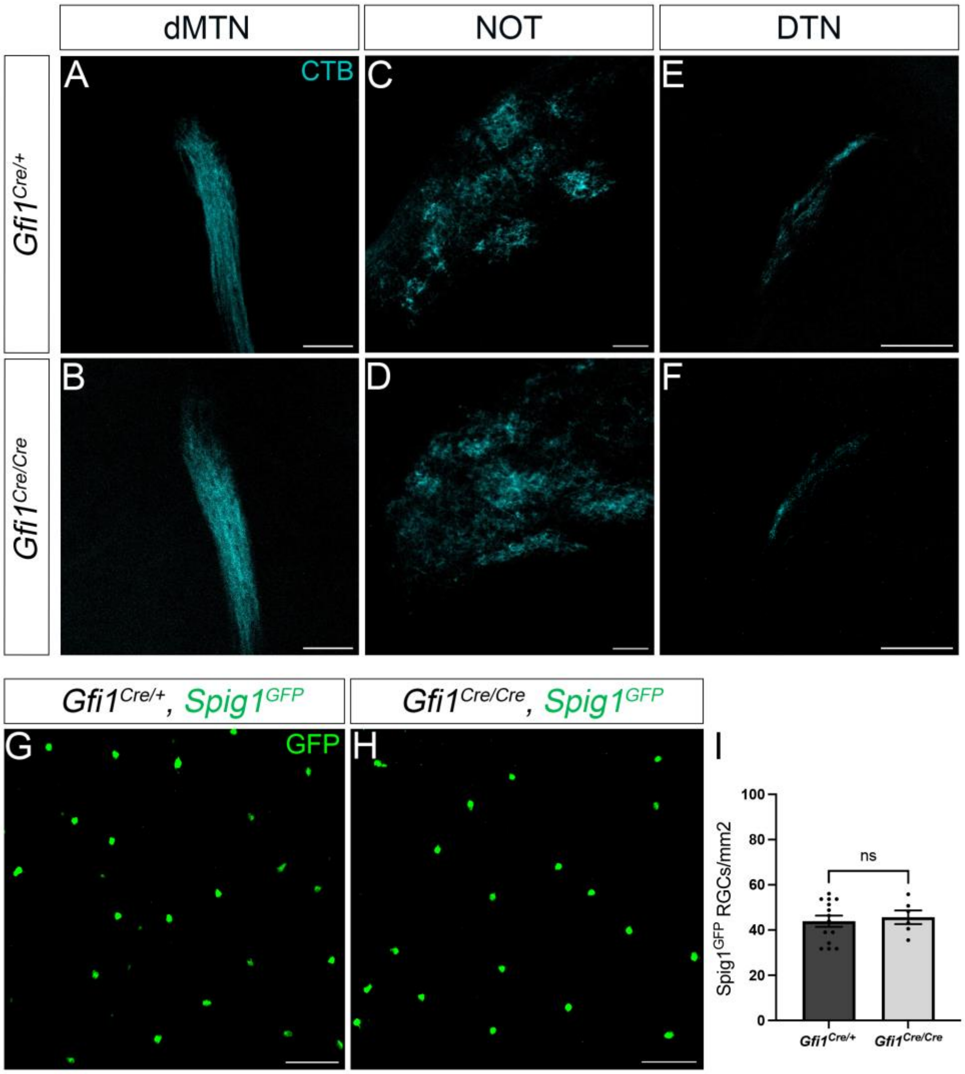


**Figure S3. Global *Gfi1* removal specifically affects D-oDSGCs within the AOS**

(**A-F**) Global Gfi1 loss does not affect innervation of other AOS nuclei including the dorsal MTN (**A, B**), NOT (**C, D**) and DTN (**E, F**). (**G-I**) The density of Spig1$^{GFP}$ U-oDSGCs remains unchanged in global *Gfi1* mutants. Data are presented as mean ± SE.

**Figure S4**

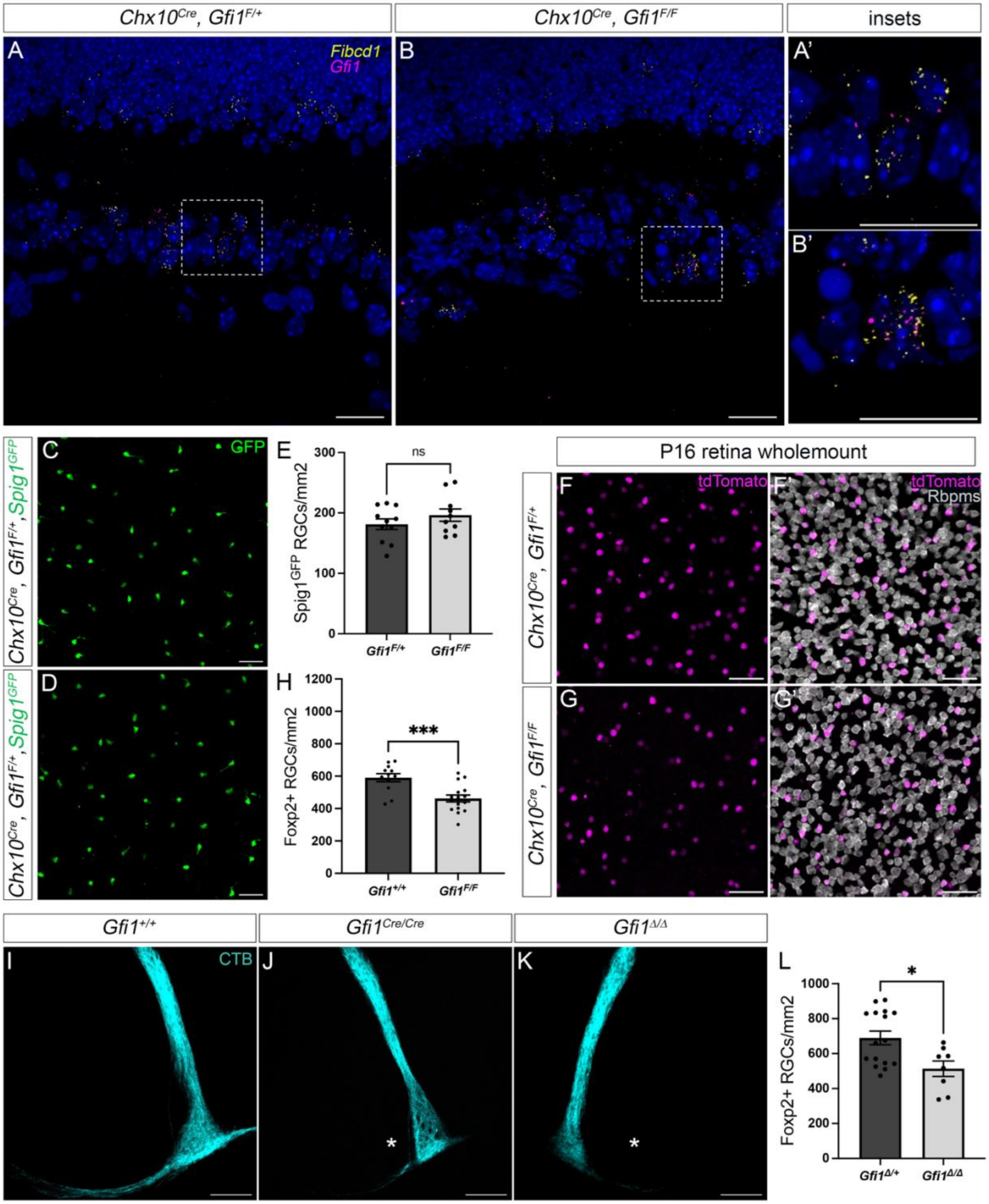

Chx10Cre, Gfi1F/+
Chx10Cre, Gfi1F/F
insets
A
Fibcd1
Gfi1
B
A'
B'
C
GFP
Chx10Cre, Gfi1F/+, Spig1GFP
D
Chx10Cre, Gfi1F/+, Spig1GFP
E
ns
Spig1GFP RGCs/mm2
Gfi1F/+
Gfi1F/F
H
***
Foxp2+ RGCs/mm2
Gfi1+/+
Gfi1F/F
P16 retina wholemount
F
tdTomato
F'
tdTomato
Rbpms
Chx10Cre, Gfi1F/+
G
G'
Chx10Cre, Gfi1F/F
Gfi1+/+
Gfi1Cre/Cre
Gfi1Δ/Δ
I
CTB
J
K
L
*
Foxp2+ RGCs/mm2
Gfi1Δ/+
Gfi1Δ/Δ

**Figure S4. Conditional *Gfi1* removal results in a loss of F-mini ONs but not D-oDSGCs**

(**A-B**) *In situ* hybridization shows *Gfi1-* and *Fibcd1*-coexpressing D-oDSGCs in P6 retina cryosections from *Chx10*$^{Cre}$*, Gfi1*$^{F/+}$ and *Chx10*$^{Cre}$*, Gfi1*$^{F/F}$ mice. (**A', B'**) Detailed views of the nuclei indicated in A and B. (**D-E**) The density of Spig1$^{GFP}$ U-oDSGCs remains unchanged in *Chx10*$^{Cre}$*, Gfi1*$^{F/F}$ retinas. (**F-H**) Wholemount retinas of *Chx10*$^{Cre}$*, Gfi1*$^{F/F}$ P16 mice show a ~25% reduction in Foxp2$^{+}$ RGCs, reflecting loss of F-mini ONs. (**I-K**) Global mutants generated from the conditional *Gfi1*$^{flox}$ allele (*Gfi1*$^{\Delta/\Delta}$) (**J**) phenocopy global *Gfi1*$^{Cre/Cre}$ mutants (**K**) and show a loss of ventral MTN innervation (indicated by asterisks) and a loss of F-mini ONs reflected by a ~25% reduction in Foxp2$^{+}$ RGCs (**L**). Data are presented as mean ± SE. *$p < 0.05$, ***$p < 0.001$.

**Figure S5**

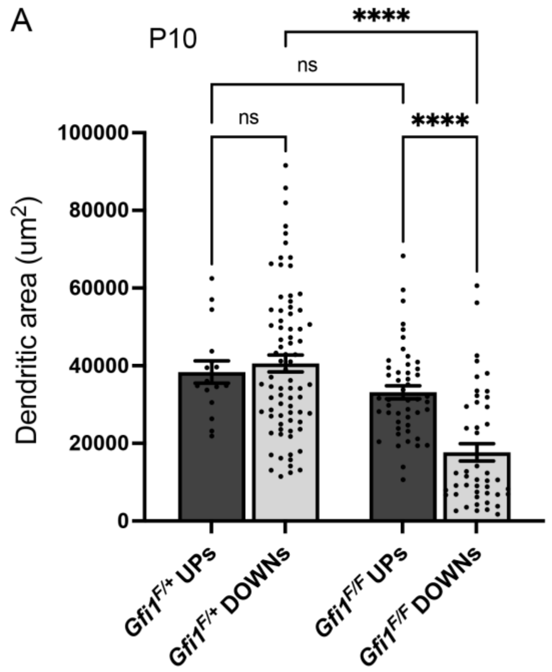


**Figure S5. The dendritic morphology defects in *Gfi1* conditional mutants persist even after DS establishment.**

(**A**) *Chx10*$^{Cre}$*, Gfi1*$^{F/F}$ mice show significantly smaller dendritic field areas of Spig1$^{GFP-}$ D-oDSGCs at P10 after the window for DS establishment which occurs ~P8. Data are presented as mean ± SE. ****$p < 0.0001$.
